# Towards Choice Engineering

**DOI:** 10.1101/2023.11.04.565653

**Authors:** Ohad Dan, Ori Plonsky, Yonatan Loewnestein

**Affiliations:** Department of Comparative Medicine, Yale University, New Haven, Connecticut; Faculty of Data and Decision Sciences, Technion – Israel Institute of Technology, Technion City, Haifa, Israel; The Edmond and Lily Safra Center for Brain Sciences, Department of Cognitive and Brain Sciences, The Alexander Silberman Institute of Life Sciences and The Federmann Center for the Study of Rationality, The Hebrew University, Jerusalem, Israel

## Abstract

Effectively shaping human and animal behavior has been of great practical and theoretical importance for millennia. Here we ask whether quantitative models of choice can be used to achieve this goal more effectively than qualitative psychological principles. We term this approach, which is motivated by the effectiveness of engineering in the natural sciences, ‘choice engineering’. To address this question, we launched an academic competition, in which the academic participants were instructed to use either quantitative models or qualitative principles to design reward schedules that maximally bias choice in a repeated, two-alternative task. We found that a choice engineering approach was the most successful method for shaping behavior in our task. This is a proof of concept that quantitative models are ripe to be used in order to engineer behavior. Finally, we show that choice engineering can be effectively used to compare models in the cognitive sciences, thus providing an alternative to the standard statistical methods of model comparison that are based on likelihood or explained variance.

## Introduction

From educating our children to persuading our friends, shaping the behavior of others is a prime goal in our society. To this end, we all use intuition and folk wisdom when interacting with fellow human beings. Salesmen and politicians often utilize established psychological principles, (e.g., anchoring, primacy, recency, etc.) to more successfully sell their products. Specifically, in behavioral economics, Nobel laureate Richard Thaler coined the term choice architecture to describe the use of qualitative psychological principles in order to influence choices without changing the choices’ objective values (Thaler, 2009).

For generations, people have been using folk physics, a qualitative intuitive understanding of the laws of nature, in order to construct tools and build buildings. However, in the last several centuries, the physical sciences have reached a level of maturity that allows us to accurately describe physical processes using mathematical equations. These equations now underlie many branches of engineering and enable the building of ever taller skyscrapers, ever faster computers, and ever more efficient cars. Motivated by the success of quantitative modeling in modern engineering, we asked whether quantitative models in the cognitive sciences can be used to efficiently engineer behavior. Quantitative modeling is prevalent when studying operant learning, the learning of association between actions and their consequences (Staddon & Cerutti, 2003), and therefore, we set out to address this question in the framework of an operant learning task.

In a typical operant learning task, a subject is instructed to repeatedly choose between two alternatives and is rewarded according to their choices (Fig. 1a). Over trials, the subject, will bias their choices in favor of the alternative that they deem more rewarding (Mongillo, Shteingart, & Loewenstein, 2014). The sequence of choice will depend on the reward schedule, that is, the mapping of choices to rewards, as well as the subject’s learning and decision-making strategies. Consider a choice engineer whose goal is to influence the subject’s sequence of choices by constructing an appropriate reward schedule. We expect that an engineer that has a deeper understanding of the strategies employed by the subject, will be able to construct a reward schedule that has a greater impact on the subject’s decision-making. This will be particularly true in complex settings.

**Figure 1.**
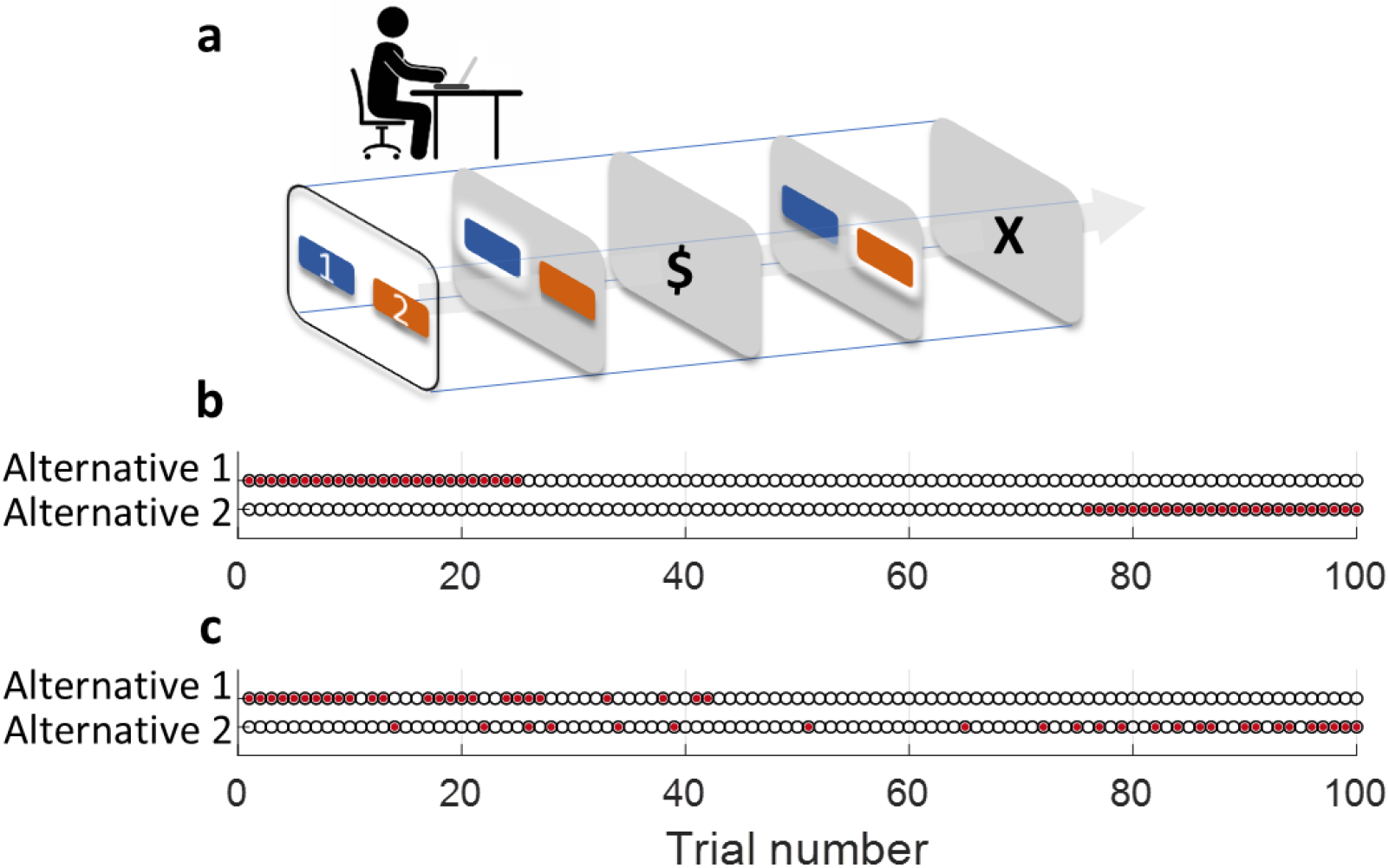
Choice engineering assignment. **a.** In the experimental task, subjects repeatedly choose between two alternatives (1 and 2). Following each choice, the subject is rewarded or not rewarded ($ or X, respectively) in accordance with a predefined binary reward schedule. Unbeknownst to participants, each of the alternatives was associated with exactly 25 rewards (red-filled circles). **b**, With the objective of maximizing bias in favor of alternative 1, primacy will favor allocating as many rewards as possible to alternative 1 at the beginning of the task. Similarly, primacy may dictate that we should defer all rewards allocated to Alternative 2 to the end of the task. **c**, A choice engineer can use a model of the subject’s learning strategy in order to construct a more effective reward schedule. The schedule depicted here is the competition winner, a reward schedule optimized for the behavioral model CATIE.

We demonstrate this idea using a specific engineering assignment: the engineer’s goal is to construct a binary reward schedule (allocate binary rewards to choices) that will maximally bias the subject’s choices in favor of choosing a predefined alternative, defined here as Alternative 1. Common sense intuition, qualitative psychological principles, and different quantitative reinforcement learning models dictate the same optimal reward schedule: the subject will be maximally biased in favor of Alternative 1 if they are rewarded whenever they choose Alternative 1, but never when they choose the other alternative, Alternative 2. In this case, an accurate model of the subject’s strategies does not seem necessary for achieving the engineer’s goal. The engineering assignment is more challenging if constraints are added to the allowed reward schedules. For concreteness, we consider the assignment of finding a reward schedule that maximally biases the subject’s choices in favor of Alternative 1 in a 100-trial session, such that the schedule assigns a binary reward to *each* of the alternatives in exactly 25 trials. Faced with this assignment, a choice architect will use qualitative psychological principles to construct an effective reward schedule. For example, primacy dictates that many rewards should be associated with Alternative 1 at the beginning of the session, whereas the association of rewards with Alternative 2 should be restricted to the end of the session (Fig. 1b). However, if the engineer possesses an accurate model of the subject’s strategies, then they can use the model to simulate the expected sequence of choices in response to different reward schedules and use these simulations to select the most effective schedule (Fig. 1c). We used this engineering assignment to ask whether quantitative models of choice can be used to engineer behavior more effectively than qualitative psychological principles and intuition.

In order to effectively engineer behavior, an accurate model of the subject’s strategies is needed. This is in the same sense that civil engineers need accurate models of the mechanics of materials when constructing large buildings and bridges. However, in the field of operant learning, there is not a uniquely-accepted model of human learning and decision-making (Mongillo et al., 2014). Rather, different models, characterized by different functional forms and parameters are used in different publications to explain learning behavior in operant tasks. Therefore, rather than committing to a specific quantitative model, we tested the effectiveness of choice engineering by launching the Choice Engineering Competition (Dan & Loewenstein, 2019). In this competition, academics were invited to propose reward schedules (with the constraints described above) with the objective of biasing the average choices of a population of subjects in favor of Alternative 1. The reward schedules could be either based on quantitative models (choice engineering) or qualitative models (choice architecture).

The effectiveness of the proposed schedules depends on the accuracy of the model that describes the subject’s learning. Therefore, the ability to engineer an effective reward schedule can be used as a measure of the accuracy of the underlying model. Engineering can thus be used as a way of comparing the “goodness” of models. Moreover, because qualitative models, as well as intuition, can also be used to construct reward schedules, this approach allowed us to compare, using the same scale, qualitative and quantitative models, a feat that cannot be achieved using standard approaches for model comparison that are based on likelihood or explained variance.

We tested the proposed reward schedules on thousands of human subjects and here we report the results of the competition. We found that a reward schedule that was based on choice engineering was comparable or better than schedules based on qualitative principles, demonstrating the potential effectiveness of choice engineering. This winning schedule was based on CATIE (Contingent Average, Trend, Inertia, and Exploration) (Plonsky & Erev, 2017), a phenomenological model of operant learning choice. Somewhat surprisingly, schedules that were based on the much more popular QL (Q-learning) model were significantly and substantially less effective in this task.

## Results

### The ideal choice engineer

There are approximately 6·10^46^ possible reward schedules consistent with the constraints of this task. The *ideal choice engineer*, an engineer that knows everything that can be known about humans’ behavior and is computationally-limitless, will select the most effective reward schedule of these possible schedules. How effective will this optimal schedule be? The maximally attainable bias depends on the learning strategies utilized by the humans. If, for example, humans employ random exploration (Fox, Dan, Elber-Dorozko, & Loewenstein, 2020) then their probability of choosing Alternative 2 will not vanish even under the optimal reward schedule. Heterogeneity between participants is another limiting factor for the maximally attainable bias (Ratcliff, Voskuilen, & McKoon, 2018; Renart & Machens, 2014). This is because a schedule that is optimal for one subject may not be that effective for another subject.

Quantifying the effectiveness of an ideal choice engineer is informative because it provides a benchmark to which the effectiveness of the reward schedules that were tested in our competition can be compared. We estimate it using two approaches. Theoretically, we searched for the optimal reward schedules assuming that subject’s learning and decision making follows specific models. Then, we used these models to estimate the bias expected from the optimal schedule. Specifically, we considered two learning strategies, QL (Sutton & Barto, 2018) and CATIE (Plonsky & Erev, 2017), two quantitative models that have been previously proposed to model choices in similar operant learning tasks. The QL model is a reinforcement learning algorithm that posits that the subject learns the expected average reward associated with each of the alternatives (also known as their values). In a trial, the subject typically chooses the alternative associated with the larger value (exploitation). Occasionally, however, the subject chooses the alternative associated with the lower value (exploration). We used an ε-softmax function to model the tradeoff between exploration and exploitation(Shteingart, Neiman, & Loewenstein, 2013). The CATIE model is more complex and describes an agent whose choices are the result of four principles. In addition to exploration and exploitation, choices in the CATIE model are also driven by inertia (repetition of the action chosen in the previous trial), and the detection of trends in the delivery of rewards. Furthermore, exploration and exploitation in CATIE are different than in many other learning models. The probability of exploration is assumed to depend on the reward prediction error: the larger the error, the higher the chances that the subject will explore (Fox et al., 2020). Importantly, rather than assuming that each alternative is associated with a static “value”, the model assumes the subject expects the outcomes to follow predictable patterns (Plonsky, Teodorescu, & Erev, 2015). Both exploitation and the computation of the reward prediction error rely on this expectation.

In the QL model, the most effective reward schedule (Fig. S1a) resulted in Alternative 1 being chosen 67.0%+/-0.1% of the trials. The CATIE model was more influenceable. The most effective reward schedule (Fig. S1b) resulted in a bias of 72.7%+/-0.1%. These numbers are only lower bounds for the performance of an ideal choice engineer with these models. This is because we could not test all possible reward schedules on these models. Rather, we used an optimization technique to identify the more effective reward schedule (Dan & Loewenstein, 2019). Therefore, it is possible (and likely) that there exist more effective schedules, although it is unlikely that they would be substantially more effective. Neither QL nor CATIE, however, are likely to fully describe humans’ learning strategies and the heterogeneity across the population. Therefore, we also addressed the question of estimating the performance of the ideal choice engineer from a different approach.

Specifically, we attempted to experimentally identify an effective reward schedule using an iterative process. We started by testing human participants using the naive reward schedule of Fig. 1b. We then calculated the empirical per-trial probability of choosing each of the targets and used these probabilities in order to construct a second, more effective reward schedule. We repeated this process three times, until the changes that we made to the reward schedule were detrimental to performance. This process resulted in the reward schedule that is depicted in Fig. S2 (top). Testing it on 549 subjects, we found that across subjects, the elicited bias was 69.0%+/-0.5%. This result is a true lower bound of the maximally attainable bias (unlike previous bounds that were based on specific models). However, this bound is less tight than the bound estimated in the simulations because we tested the subjects using a relatively small number of reward schedules.

The fact that we found a reward schedule that induces a bias in human subjects that is substantially and significantly larger than the maximally attainable bias of the QL model is an indication that QL does not describe humans’ strategies well. More on that later.

### Choice engineering in the competition

In the competition, we tested the effectiveness of 11 different reward schedules on thousands of human subjects, each tested on a single schedule. Of the 11 schedules, 7 were based on quantitative models, and 4 were based on qualitative principles (see Table S1 and Fig. S2). The fraction of trials in which Alternative 1 was chosen in a reward schedule, averaged over all trials and over all subjects is depicted in Fig. 2. We found that there was a substantial and significant difference in the efficacy of the different schedules. While the least effective schedule was not significantly different from chance (50.7%+/-1.8%, p=0.72), the best performing reward schedule yielded significant bias of 64.3%+/-0.5% (p<0.001). This reward schedule was submitted by a choice engineer who optimized the reward schedule based on CATIE. Second, came a reward schedule that was submitted by a team of choice architects that combined primacy with strategic considerations to design the reward schedule (Table S1). The average bias of that model was 64.1%+/-0.6% not significantly different from that of the first place (p=0.84, df=1131). The schedule that came third, also constructed by a choice architect, attempted to dynamically manipulate the level of exploration. It achieved a bias of 62.7%+/-0.6%, significantly lower than the winning schedule (p=0.037, df=1200). All other schedules achieved a significantly lower bias compared to the winning schedule (all p<0.004).

**Figure 2.**
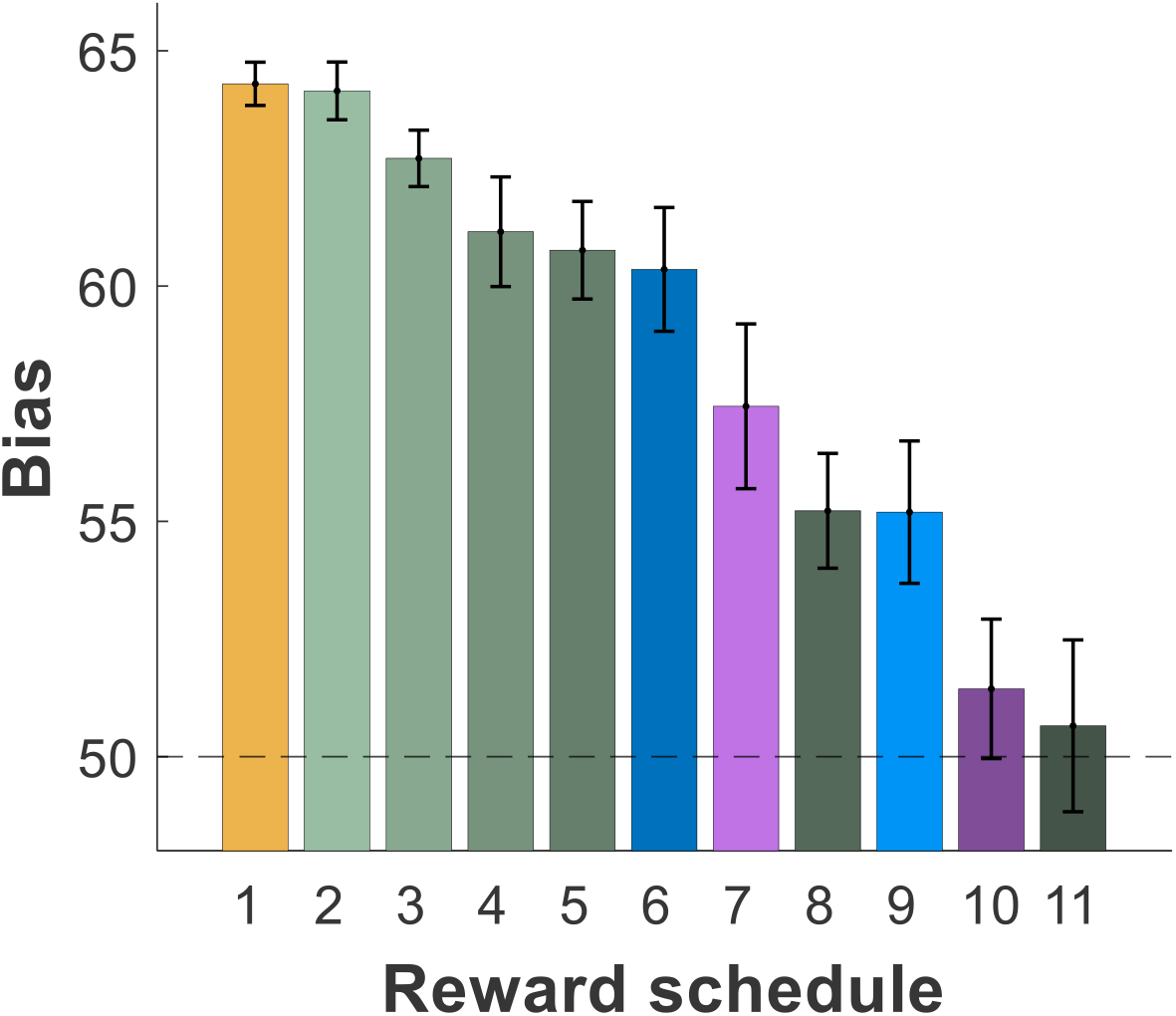
Competition results. The bias, average proportion of choices of Alternative 1 for the different reward schedules (Fig. S2 and Table S1). The winner of the competition (schedule 1, orange) was designed by a choice engineer that utilized the CATIE model, achieving an average bias of 64.3%. Noteworthy are also the results of schedules optimized using a QL model with a single set of parameters (“homogeneous”, schedules 7 and 9) or an ensemble of parameters (“heterogeneous”, schedules 6 and 10). The parameters of the models used previous studies (‘lab’, schedules 7 and 10) or data released for the competition (‘online’, schedules 6 and 9). Error bars are standard error of the means. Note that, to optimize the chances of identifying the better schedules, a different number of subjects was used for each schedule. Therefore, the more successful scheduled were tested on more subjects (Table S1).

Specifically, four different reward schedules were engineered based on the popular QL model. These models differed in their parameters and therefore yielded four different schedules. In two models, the parameters were estimated from a previous study (Erev et al., 2010; Shteingart et al., 2013) that was conducted on humans in the laboratory. The parameters of the two other models were estimated from a dataset of online operant learning experiments that we released prior to the competition. We denote these models as ‘lab’ and ‘online’, respectively. Moreover, in two models, the parameters were fitted assuming that a single set of parameters describes all subjects. In the two other models, the idiosyncratic parameters of each subject were estimated. Remarkably, all four schedules fared considerably worse than the CATIE-based schedule, a single schedule whose parameters were based on previous laboratory studies and were not fitted to online dataset.

### The CATIE model and the competition

The success of the CATIE-based reward schedule suggests that CATIE is a good description of humans’ learning and decision-making strategies. Nevertheless, its performance of 64.3%+/-0.5% was substantially lower than the expected performance of an ideal choice engineer given the CATIE model, 72.4%+/-0.1%. This lower performance was not due to a failure of the winning engineer in finding an optimal schedule for the CATIE model, as simulating the winning schedule using the CATIE model resulted in a bias of 71.6%+/-0.1%, comparable to that of our optimal schedule. These results indicate that the failure of the winning schedule is due to an inaccurate description of humans’ learning behavior.

To further quantify this inaccuracy, we simulated each of the schedules using the CATIE model and compared the mean biases of these simulations to the empirically-measured mean biases (Fig. 3a). Qualitatively, the simulations exhibited a similar dependence of the bias on the schedule but there were still some differences (see also below). One possible cause for the discrepancies between the simulations and humans’ behavior could have been that humans are substantially more heterogeneous in their learning strategy (Berry, Jagust, & Hsu, 2018; Laquitaine, Piron, Abellanas, Loewenstein, & Boraud, 2013; Lebovich, Darshan, Lavi, Hansel, & Loewenstein, 2019; Peters & Büchel, 2011; Wyart & Koechlin, 2016) than is accounted for by the model. To test this possibility, we used the fact that a large number of participants were tested on the same reward schedule and we can quantify their heterogeneity. For example, while averaged over subjects, Alternative 1 was chosen in 64.3%+/-0.5% of the trials when tested using the winning schedule, some subjects chose it in more than 90% of the trials whereas others chose it in less than 50% of the trials (see Fig. S3). The standard deviation of the bias, 11.2%+/-0.04%, is a measure of heterogeneity in their performance. This heterogeneity, however, does not necessarily imply that subjects employed different strategies. The reason is that the decision-making strategies can incorporate stochasticity, which can account for variability in their choices (for example, random exploration is a component of both QL and CATIE models). Therefore, we compared the standard deviation of the bias measured over the human subjects to that predicted by the CATIE model. We found that for the winning schedule, the experimentally-observed standard deviation was 11.2%+/-0.04%, was only slightly greater than predicted by the CATIE model 10.0%+/-0.02. This result suggests that heterogeneity between subjects’ strategies was not substantial in this task.

**Fig 3.**
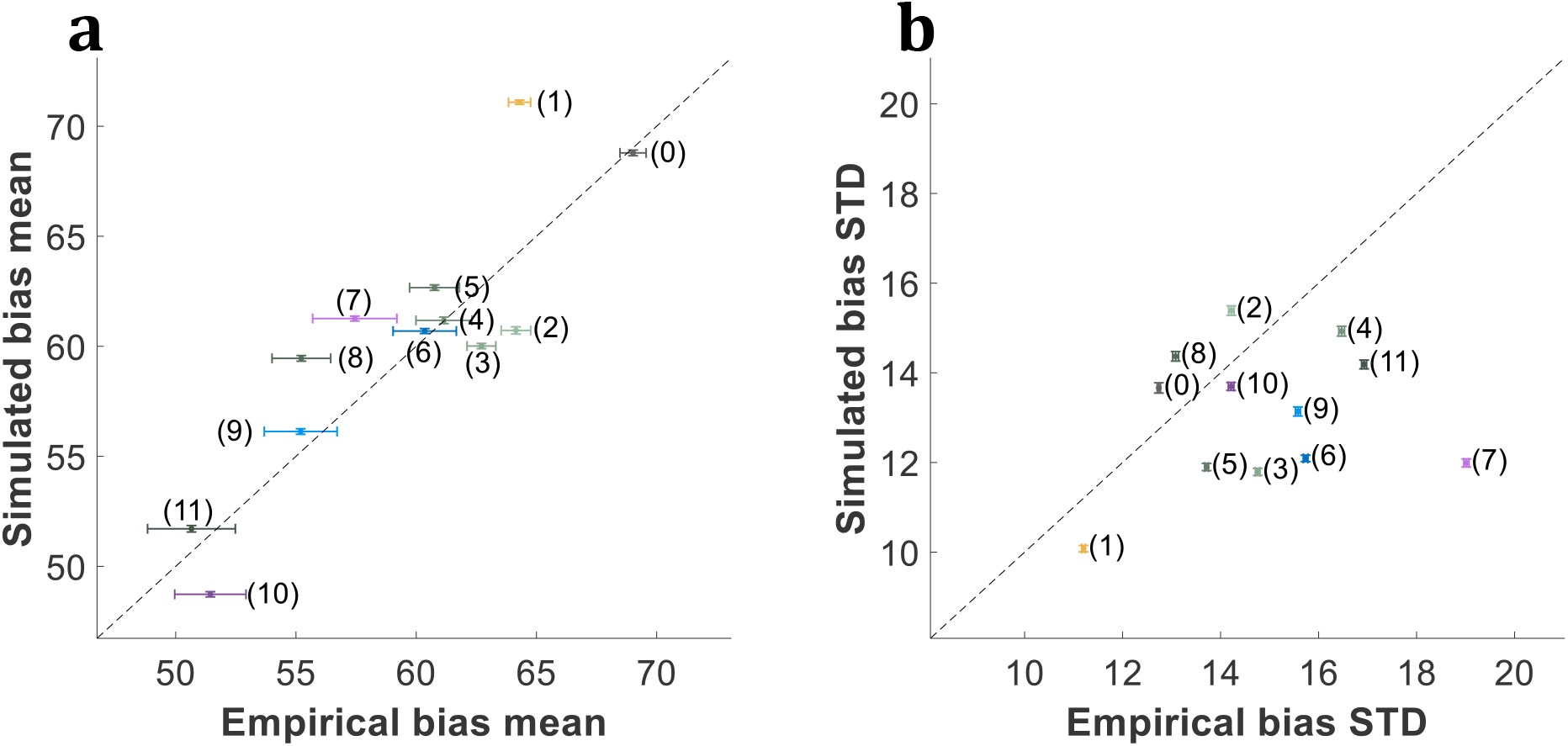
Predicted versus experimentally-measured bias. **a,** The mean bias for each of the reward schedules of Fig. S2, as predicted by the CATIE model, versus the mean experimentally-measured bias. **b,** The same as **a**, for the standard deviation (over the subjects). The simulated standard deviation was computed by averaging over 1,000 simulations, each with the same number of participants as in the competition. Error bars are standard error of these means. Dashed line is the diagonal.

To further test it, we compared the experimentally-observed standard deviations of the bias of the human subjects to that predicted by the CATIE model for all schedules (Fig. 3b, see also Fig. S3) and found that humans were only marginally more heterogeneous than predicted by the model (the distribution of biases was wider for 3/12 schedules, p=0.07). These results further indicate that heterogeneity between subjects did not play an important role in this task. Consequently, we conclude that a substantial contributor to the difference between the performance of the ideal engineer and the winning engineer is the fact that CATIE does not fully capture humans’ learning strategy.

### Choice engineering and standard methods of model comparison

In the competition, a schedule that was based on CATIE fared better than schedules that were based on QL. This raises the question of whether CATIE also *describes* humans’ choices better than QL. To address this question, we considered three measures of model description:

a. The objective of the participants in the competition was to bias choices in favor of Alternative 1. Therefore, a natural measure of performance of the different models is their ability to *predict* the bias from the schedule. To this end, we simulated the four versions of the QL model and the CATIE model and compared the models’ bias predictions to those experimentally measured for each of the tested reward schedules (Fig. S4). One measure of the agreement between the model and the experimental results is the correlation coefficient *r* of the predicted bias with the experimentally-measured bias. We found that the correlation coefficient of the CATIE model *r*=0.88 was substantially larger than that of all other models (Fig. 4a).
b. Another way of quantifying the accuracy of a model is to compare the probabilities *p* that it assigns to the subjects’ choices, where these probabilities are based on the model parameter and past choices (Materials and Methods). The larger *p*, averaged over trials and participants (E[*p*]), the more accurate is the model. Because the parameters of the models were determined independently prior to the experiment, there was no need to correct for the number of parameters. As depicted in Figure 4b, CATIE outperformed the other models in this measure with E[*p*]=61.9%, a number that is marginally higher than E[*p*]=61.8%, the homogeneous online QL model that came second (p=0.054).
c. Finally, a standard way of comparing models is based on the likelihood of the data given the model. Ranking is done by comparing E[log(*p*)], where the larger the number (closer to zero), the more accurate the model (Fig. 4c). In this analysis, QL, heterogeneous, online obtained the highest likelihood, whereas the CATIE model came only third.

**Figure 4.**
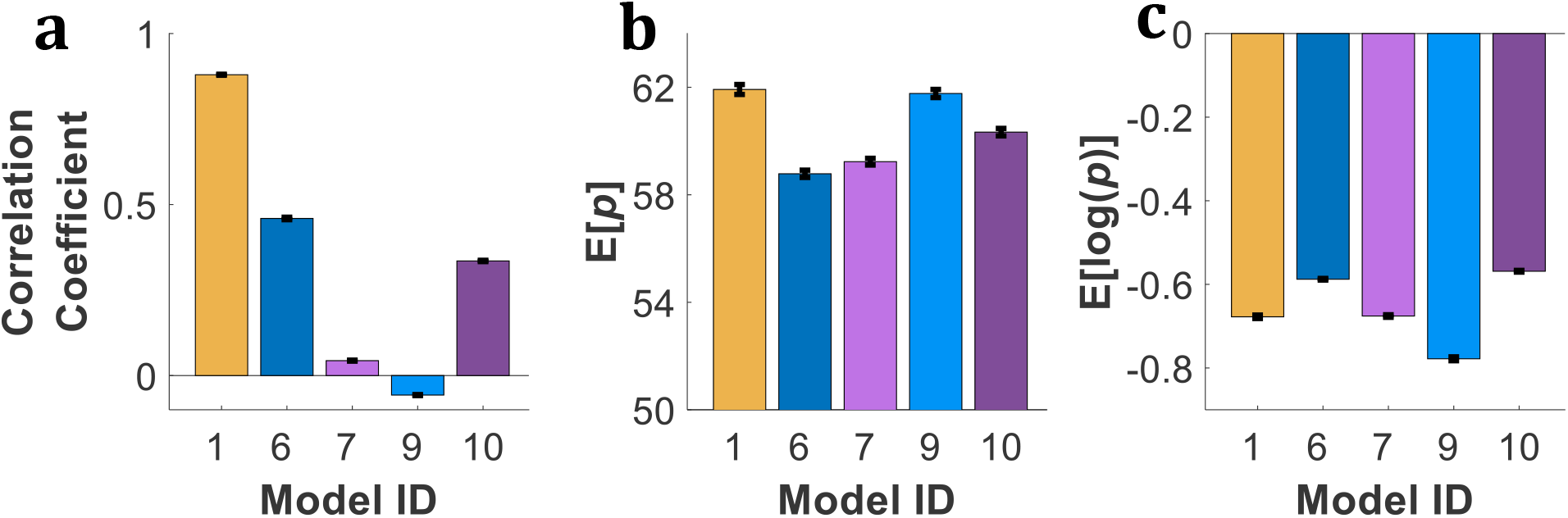
Model comparison. We used three criteria to compare the abilities of four versions of QL and CATIE to describe subjects’ choices. **a**, Correlation of predicted and empirical bias over the different schedules; **b**, Average choice probability, E[*p*] **c**, Log-likelihood, E[log(*p*)]. While CATIE (orange bar) was the best model with respect to average choice probability and predicted bias, the heterogeneous online QL model (light red) was superior with respect to log-likelihood.

Worthwhile noting that while 11 schedules were tested in the competition, only five models were compared with respect to their abilities to describe the data. This is because while qualitative models can be used to construct successful reward schedules (‘architect’), only quantitative models can be used to compare models by their ability to quantitatively describe behavior.

## Discussion

In this paper we demonstrated that quantitative models of choice are capable of efficiently engineering behavior. These results are a proof of principle of the potential of choice engineering. Faced with an objective function and a quantitative model of humans’ learning strategy, the winning choice engineer successfully designed an effective reward schedule using standard optimization techniques. This is an approach that can be automatically applied to other objective functions and constraints. By contrast, the choice architect approach that is based on human expertise and intuition cannot be easily automated and any change in the objectives and constraints would require the laborious design of a new schedule.

### Operant learning

In the last decades, reinforcement learning algorithms have dominated the field of operant learning. These algorithms have been used to model human and animal learning behaviors and effort has been made to identify the brain regions, neurons and neurotransmitters that implement them (Botvinick, Wang, Dabney, Miller, & Kurth-Nelson, 2020). This development has been motivated in part by the proven ability of these algorithms to learn complicated tasks, from motor control tasks to playing games without prior knowledge (Mousavi, Schukat, & Howley, 2018). The QL algorithm is a most successful example. In machine learning, it converges (under certain conditions) to the optimal solution. In the cognitive sciences, it has been widely used to model the behavior of animals and humans. In neuroscience, its variables have been linked to neural activity in specific neurons in specific brain regions. In view of these, the success of CATIE is surprising. Particularly remarkable is the fact that the parameters of this model were not fitted to this particular competition. Rather, they were taken from previous experiments. The CATIE model is a phenomenological model that has not been derived from first principles (e.g., optimality) and has not been proven effective in the field of machine learning. Its success, therefore, may indicate that caution should be exercised when using reinforcement learning algorithms to model human learning strategies.

### Model comparison

Model comparison is becoming increasingly popular in the cognitive sciences. Methods that are based on maximum likelihood are popular because they have a Bayesian interpretation. If we know that behavior was generated by one of the models and a-priori they are equally likely, the model associated with the maximum likelihood is the one most likely to be the correct one. However, if none of the models is exact (as is always the case), there is no clear interpretation to higher likelihood of one model compared to another. Therefore, there is no unique method for model comparison (see also Figure 4). Indeed, different descriptions of behavior favored different models. The ability of the model to engineer behavior is an alternative approach to model comparison. This approach is also not unique as there is no general rule that dictates the proper task for engineering. In fact, in the academic competition (Dan & Loewenstein, 2019) we offered two tracks. In the “static” track, described in this paper, academic participants proposed a full reward schedule, as is described in this paper. In the “dynamic” track they could use a more sophisticated approach and dynamically allocate, in each trial, the reward(s) of the next trial. Regrettably, only choice architects participated in the dynamic track and therefore, we were unable to use it in order to compare choice architecture to choice engineering (see Table S2 in the Supporting Information section). The major advantage of the engineering approach over the standard methods for model comparison is that it allows the comparison of quantitative models with qualitative principles.

### Social and ethical implications of choice engineering

Influencing humans’ preferences and choices is a major objective in our society, and scientific research has been harnessed to achieve this goal (Fennis & Stroebe, 2015). Specifically, psychological principles have been used to construct effective advertising and political campaigns for over a century. More recently, large data and statistical methods are used, especially in online advertising, to achieve similar goals (Hofacker, Malthouse, & Sultan, 2016; Lee & Cho, 2019). Choice engineering can be used as an additional tool in this toolkit. Its main advantage is that it does not need large data. Given a model, the objectives can be achieved almost automatically by solving an optimization problem. However, while quantitative models for operant learning exist, it is not clear how this approach can be applied to biasing humans towards buying one brand of soft drink over another, or voting for one political candidate and not another. One limitation of the winning CATIE model, compared with QL, is in generalization to other operant problems. While QL can be directly applied to a large family of decision problems known as Markov Decision Problems, the generalization of CATIE to this family of decision problems is unclear. Despite these limitations, we predict that choice engineering will become a common tool in coming years. Therefore, the social and ethical implications of choice engineering should be considered.

## Materials and Methods

The study was approved by the Hebrew University Committee for the Use of Human Subjects in Research and all participants provided informed consent. Recruitment was based on the online labor market Amazon Mechanical Turk. Data for the static track were collected from 3,386 subjects. Subjects received a fixed participation fee of $0.4, and a bonus of additional 1¢ for every obtained reward. We considered all subjects who completed the experiment (100 choices). We excluded from the analysis 54 subjects (1.6%) that chose one of the alternatives less than 5 times and are therefore likely to have ignored the reward schedule. Adding them to the analysis does not change the ranking of the first five schedules. We did not collect any demographic information about the subjects.

Competition rules are described in detail in (Dan & Loewenstein, 2019) and are available on the competition website at http://decision-making-lab.com/competition/index.html. Briefly, we published an open call for participation in the competition (Dan & Loewenstein, 2019) and accepted all valid submissions. The goal of the participants in the competitions was to generate the reward schedule that is the most effective in biasing human subjects’ choices in favor of Alternative 1, as described in the Introduction. We also invited participants to contribute a dynamic reward schedule in which in each trial, the rewards are allocated based on past choices. However, only choice architects participated in the dynamic track and therefore, we were unable to use it in order to compare choice architecture to choice engineering (see Table S2 in the Supporting Information section).

To identify the most effective schedule, we tested the schedules on human subjects. As described in (Dan & Loewenstein, 2019), we utilized an adaptive statistical method that allocates a larger number of participants to schedules with better performance (Supplementary text).

The QL and CATIE models, as well as the schedule optimization algorithm are described in (Dan & Loewenstein, 2019).

For the model comparison of Fig. 4, we predicted the choice of each subject in each trial *t* based on all her choices in preceding trials 1 to *t*-1 and the model. In heterogeneous models, we fit a parameter distribution over the population and weighted predictions based on each set’s likelihood of generating the observed choices. Specifically, for each subject and for each trial *t*, we calculated the likelihood that her choices in trials 1 to *t*-1 were generated by each set of parameters, and then used this to weight their choice predictions in trial *t*.

## Acknowledgments

We would like to thank Lea Kaplan for technical assistance, Peter Dayan, Amir Dezfouli, Ido Erev, Yosef Rinott and Jon Roiser for helpful discussions and all the participants in the competition. This work was supported by the Israel Science Foundation (Y.L.), the Gatsby Charitable Foundation (Y.L.) and the David and Inez Myers Chair in Neural Computation (Y.L).

## Author contributions

OD and YL analyzed the data and wrote the manuscript. OP contributed the winning schedule.

## Data availability

Data files can be found in the competition website: https://sites.google.com/view/cec19/home.

### Supplementary

**Table S1.**
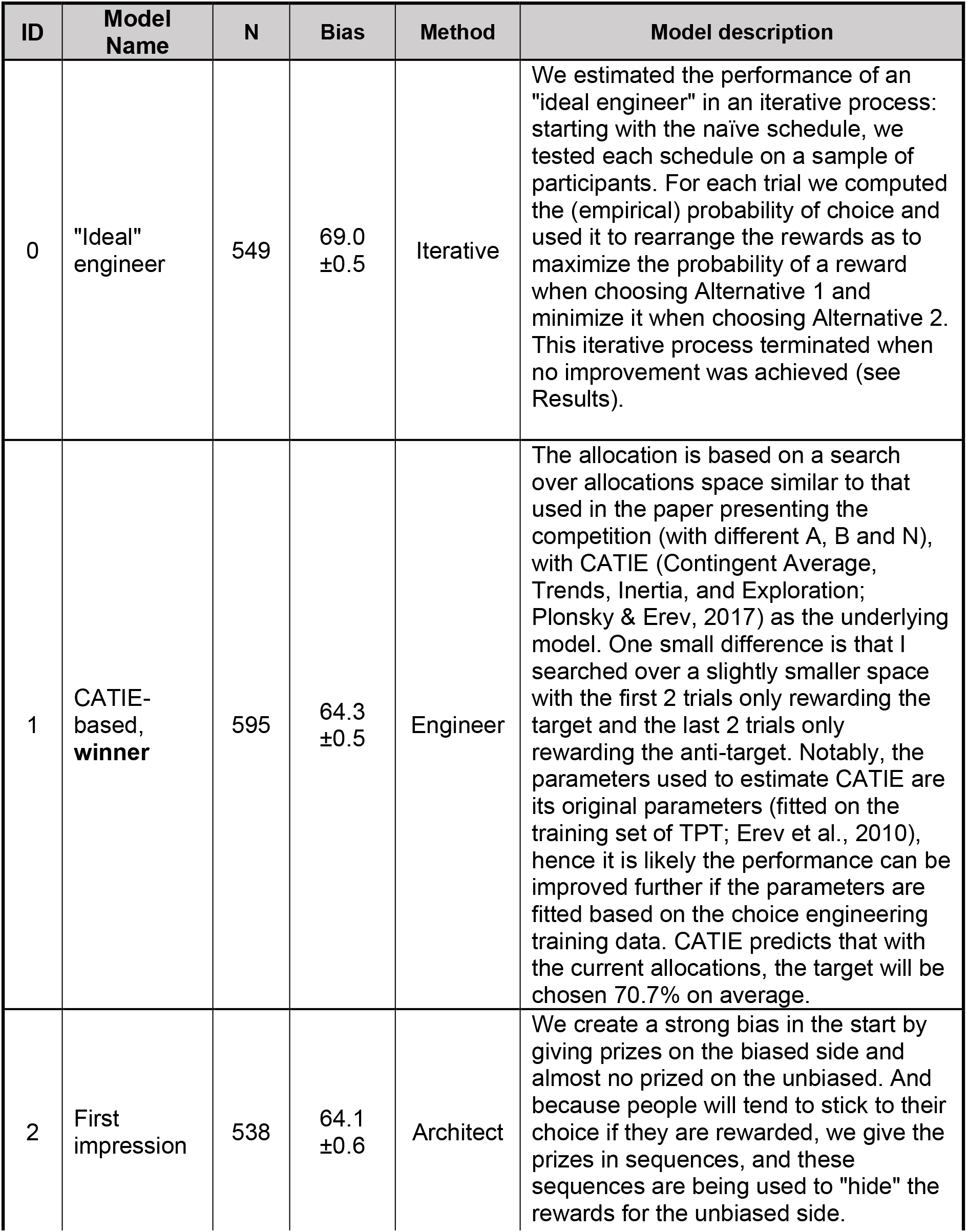
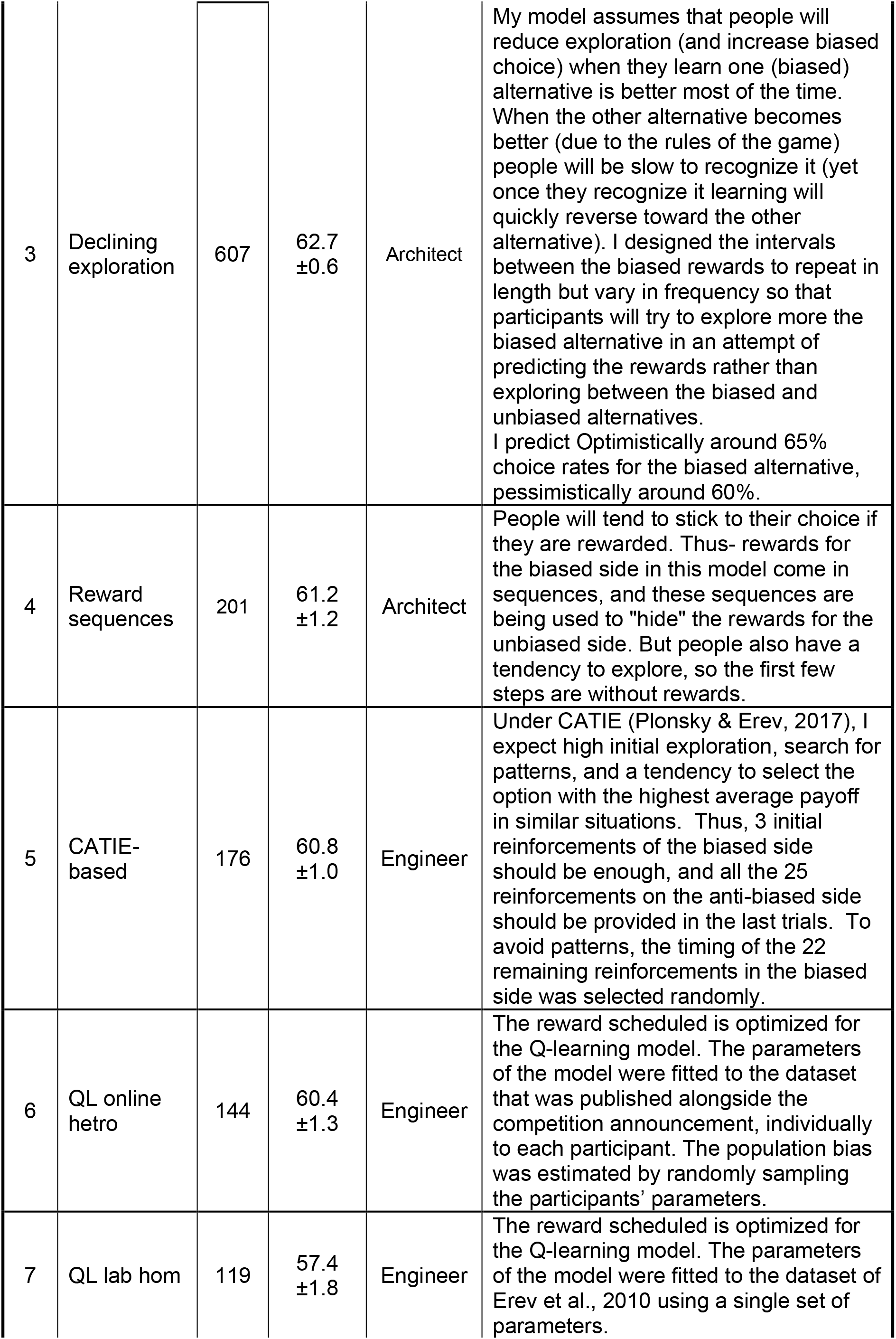
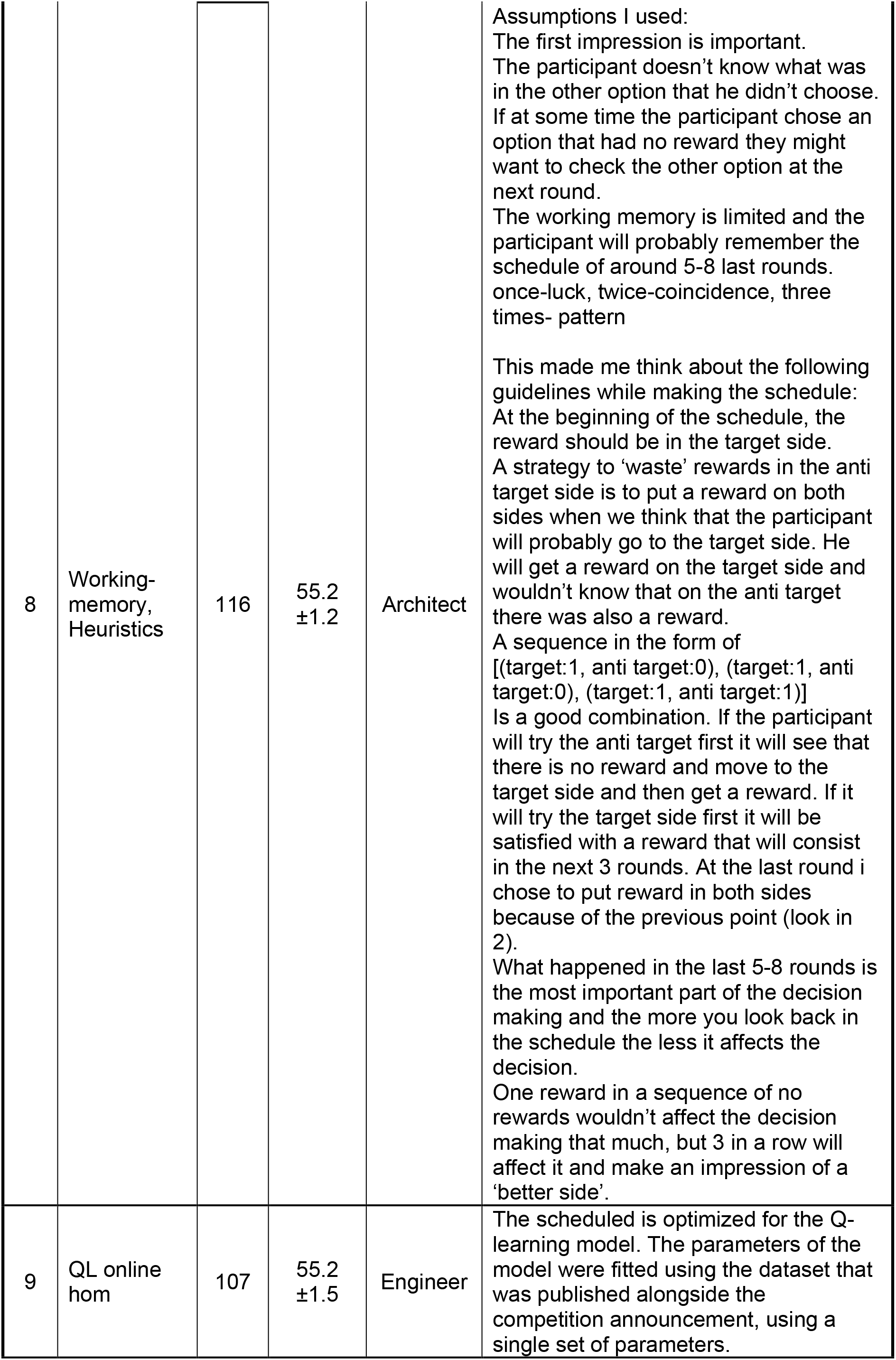
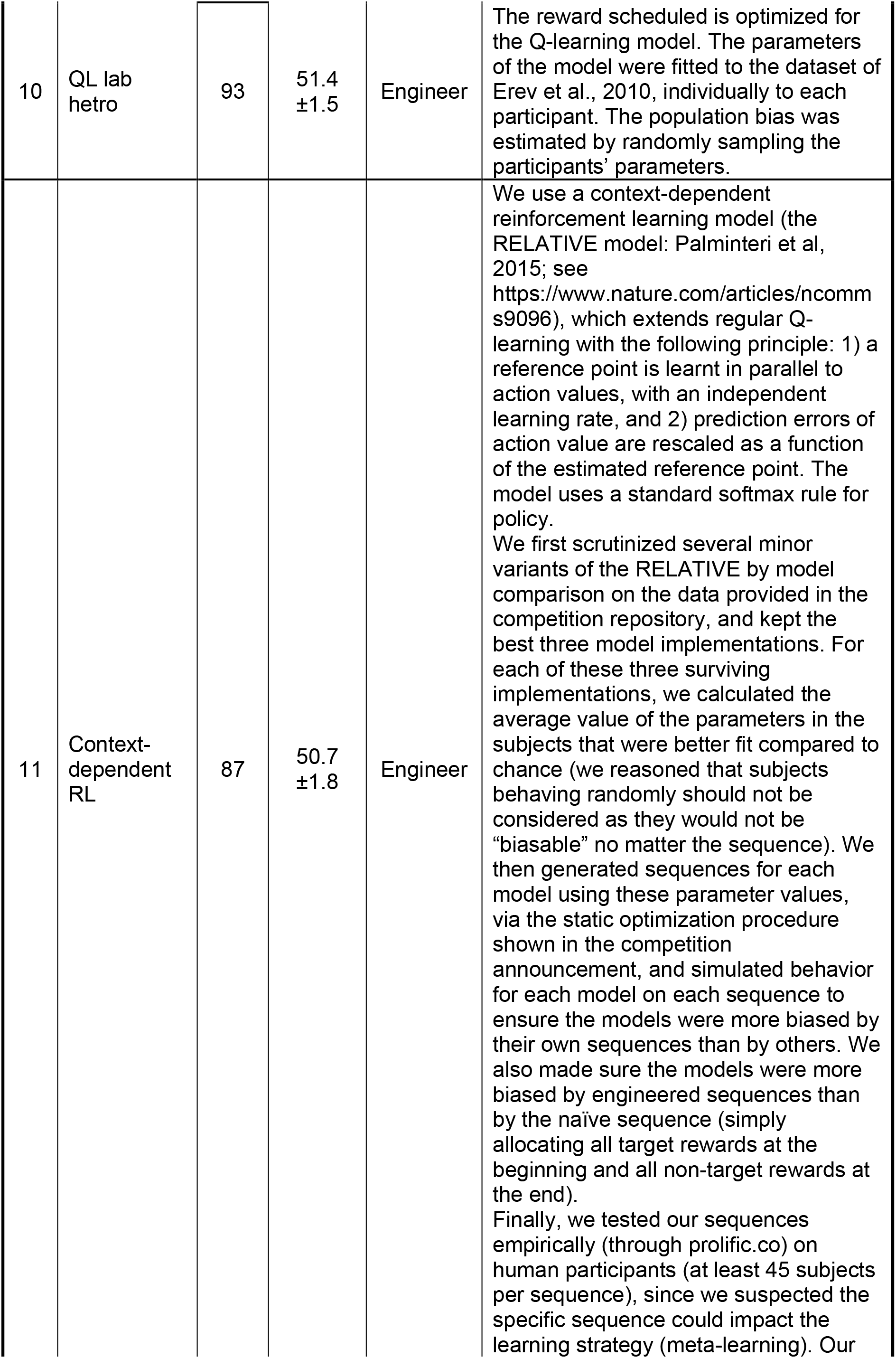
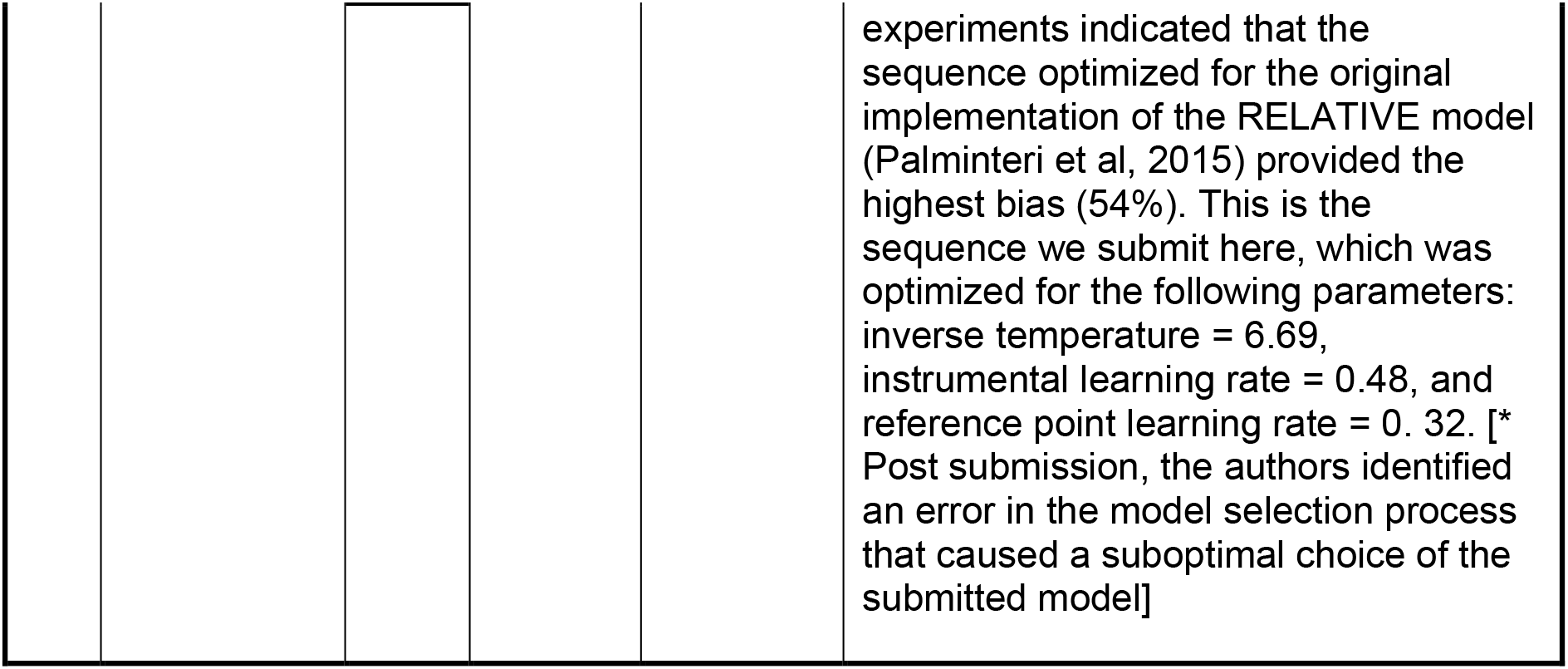
Reward schedules. The table shows the details of the 11 reward schedules tested in the competition. Schedule IDs (first column) correspond to the schedule IDs shown in Fig S1. We asked participants to report whether they used qualitative or quantitative approaches in designing the reward schedules, making them choice architects or choice engineers as indicated in the third column. In addition, participants provided a short explanation of the method they used to create the schedule. An additional reward schedule (Model ID: 1) was designed by an iterative process to estimate the performance of an “ideal engineer” (see Results).

| ID | Model Name | N | Bias | Method | Model description |
| --- | --- | --- | --- | --- | --- |
| 0 | "Ideal" engineer | 549 | 69.0<br>±0.5 | Iterative | We estimated the performance of an "ideal engineer" in an iterative process: starting with the naïve schedule, we tested each schedule on a sample of participants. For each trial we computed the (empirical) probability of choice and used it to rearrange the rewards as to maximize the probability of a reward when choosing Alternative 1 and minimize it when choosing Alternative 2. This iterative process terminated when no improvement was achieved (see Results). |
| 1 | CATIE-based, **winner** | 595 | 64.3<br>±0.5 | Engineer | The allocation is based on a search over allocations space similar to that used in the paper presenting the competition (with different A, B and N), with CATIE (Contingent Average, Trends, Inertia, and Exploration; Plonsky & Erev, 2017) as the underlying model. One small difference is that I searched over a slightly smaller space with the first 2 trials only rewarding the target and the last 2 trials only rewarding the anti-target. Notably, the parameters used to estimate CATIE are its original parameters (fitted on the training set of TPT; Erev et al., 2010), hence it is likely the performance can be improved further if the parameters are fitted based on the choice engineering training data. CATIE predicts that with the current allocations, the target will be chosen 70.7% on average. |
| 2 | First impression | 538 | 64.1<br>±0.6 | Architect | We create a strong bias in the start by giving prizes on the biased side and almost no prizes on the unbiased. And because people will tend to stick to their choice if they are rewarded, we give the prizes in sequences, and these sequences are being used to "hide" the rewards for the unbiased side. |
| 3 | Declining exploration | 607 | 62.7<br>±0.6 | Architect | My model assumes that people will reduce exploration (and increase biased choice) when they learn one (biased) alternative is better most of the time. When the other alternative becomes better (due to the rules of the game) people will be slow to recognize it (yet once they recognize it learning will quickly reverse toward the other alternative). I designed the intervals between the biased rewards to repeat in length but vary in frequency so that participants will try to explore more the biased alternative in an attempt of predicting the rewards rather than exploring between the biased and unbiased alternatives.<br>I predict Optimistically around 65% choice rates for the biased alternative, pessimistically around 60%. |
| 4 | Reward sequences | 201 | 61.2<br>±1.2 | Architect | People will tend to stick to their choice if they are rewarded. Thus- rewards for the biased side in this model come in sequences, and these sequences are being used to "hide" the rewards for the unbiased side. But people also have a tendency to explore, so the first few steps are without rewards. |
| 5 | CATIE-based | 176 | 60.8<br>±1.0 | Engineer | Under CATIE (Plonsky & Erev, 2017), I expect high initial exploration, search for patterns, and a tendency to select the option with the highest average payoff in similar situations. Thus, 3 initial reinforcements of the biased side should be enough, and all the 25 reinforcements on the anti-biased side should be provided in the last trials. To avoid patterns, the timing of the 22 remaining reinforcements in the biased side was selected randomly. |
| 6 | QL online hetro | 144 | 60.4<br>±1.3 | Engineer | The reward scheduled is optimized for the Q-learning model. The parameters of the model were fitted to the dataset that was published alongside the competition announcement, individually to each participant. The population bias was estimated by randomly sampling the participants' parameters. |
| 7 | QL lab hom | 119 | 57.4<br>±1.8 | Engineer | The reward scheduled is optimized for the Q-learning model. The parameters of the model were fitted to the dataset of Erev et al., 2010 using a single set of parameters. |
| 8 | Working-memory, Heuristics | 116 | 55.2<br>±1.2 | Architect | Assumptions I used:<br>The first impression is important.<br>The participant doesn't know what was in the other option that he didn't choose.<br>If at some time the participant chose an option that had no reward they might want to check the other option at the next round.<br>The working memory is limited and the participant will probably remember the schedule of around 5-8 last rounds.<br>once-luck, twice-coincidence, three times- pattern<br>This made me think about the following guidelines while making the schedule:<br>At the beginning of the schedule, the reward should be in the target side.<br>A strategy to 'waste' rewards in the anti target side is to put a reward on both sides when we think that the participant will probably go to the target side. He will get a reward on the target side and wouldn't know that on the anti target there was also a reward.<br>A sequence in the form of [(target:1, anti target:0), (target:1, anti target:0), (target:1, anti target:1)]<br>Is a good combination. If the participant will try the anti target first it will see that there is no reward and move to the target side and then get a reward. If it will try the target side first it will be satisfied with a reward that will consist in the next 3 rounds. At the last round i chose to put reward in both sides because of the previous point (look in 2).<br>What happened in the last 5-8 rounds is the most important part of the decision making and the more you look back in the schedule the less it affects the decision.<br>One reward in a sequence of no rewards wouldn't affect the decision making that much, but 3 in a row will affect it and make an impression of a 'better side'. |
| 9 | QL online hom | 107 | 55.2<br>±1.5 | Engineer | The scheduled is optimized for the Q-learning model. The parameters of the model were fitted using the dataset that was published alongside the competition announcement, using a single set of parameters. |
| 10 | QL lab hetro | 93 | 51.4<br>±1.5 | Engineer | The reward scheduled is optimized for the Q-learning model. The parameters of the model were fitted to the dataset of Erev et al., 2010, individually to each participant. The population bias was estimated by randomly sampling the participants' parameters. |
| 11 | Context-dependent RL | 87 | 50.7<br>±1.8 | Engineer | We use a context-dependent reinforcement learning model (the RELATIVE model: Palminteri et al, 2015; see https://www.nature.com/articles/ncomms9096), which extends regular Q-learning with the following principle: 1) a reference point is learnt in parallel to action values, with an independent learning rate, and 2) prediction errors of action value are rescaled as a function of the estimated reference point. The model uses a standard softmax rule for policy.<br>We first scrutinized several minor variants of the RELATIVE by model comparison on the data provided in the competition repository, and kept the best three model implementations. For each of these three surviving implementations, we calculated the average value of the parameters in the subjects that were better fit compared to chance (we reasoned that subjects behaving randomly should not be considered as they would not be "biasable" no matter the sequence). We then generated sequences for each model using these parameter values, via the static optimization procedure shown in the competition announcement, and simulated behavior for each model on each sequence to ensure the models were more biased by their own sequences than by others. We also made sure the models were more biased by engineered sequences than by the naïve sequence (simply allocating all target rewards at the beginning and all non-target rewards at the end).<br>Finally, we tested our sequences empirically (through prolific.co) on human participants (at least 45 subjects per sequence), since we suspected the specific sequence could impact the learning strategy (meta-learning). Our |

**Table S1. Reward schedules.** The table shows the details of the 11 reward schedules tested in the competition.
|  |  |  |  |  |
| --- | --- | --- | --- | --- |
|  |  |  |  | experiments indicated that the sequence optimized for the original implementation of the RELATIVE model (Palminteri et al, 2015) provided the highest bias (54%). This is the sequence we submit here, which was optimized for the following parameters: inverse temperature = 6.69, instrumental learning rate = 0.48, and reference point learning rate = 0.32. [* Post submission, the authors identified an error in the model selection process that caused a suboptimal choice of the submitted model] |

**Fig S1.**
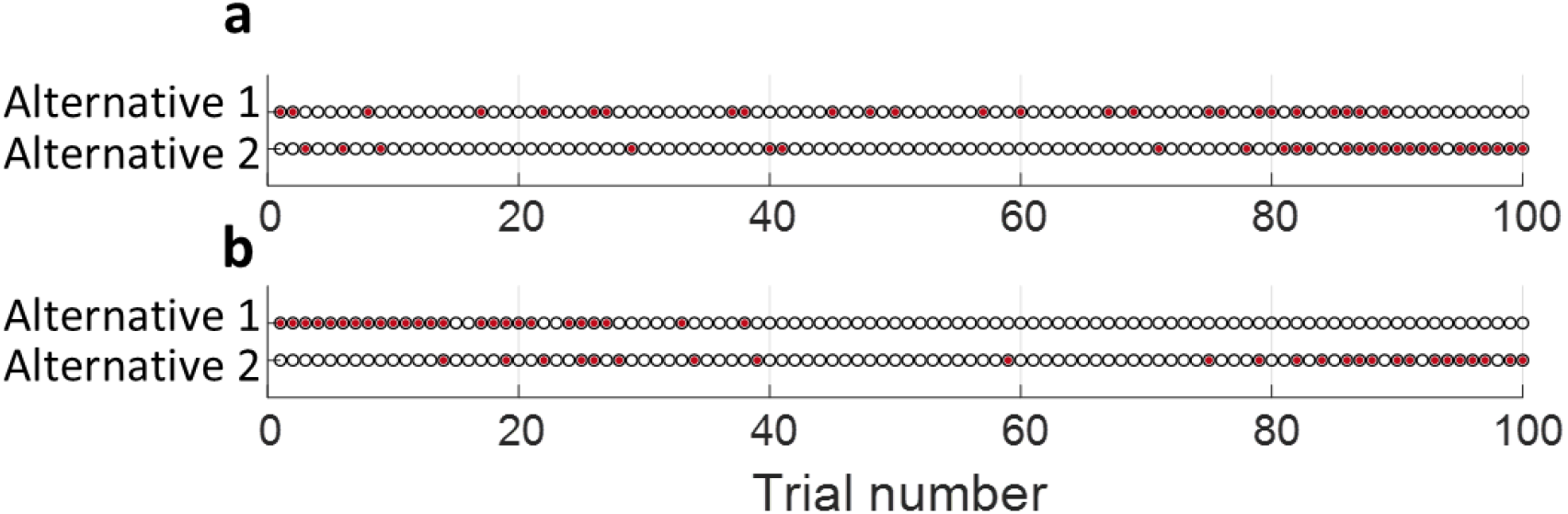
Optimized reward schedules. **a** and **b,** Reward schedules, optimized for the QL and CATIE models, respectively.

**Fig S2.**
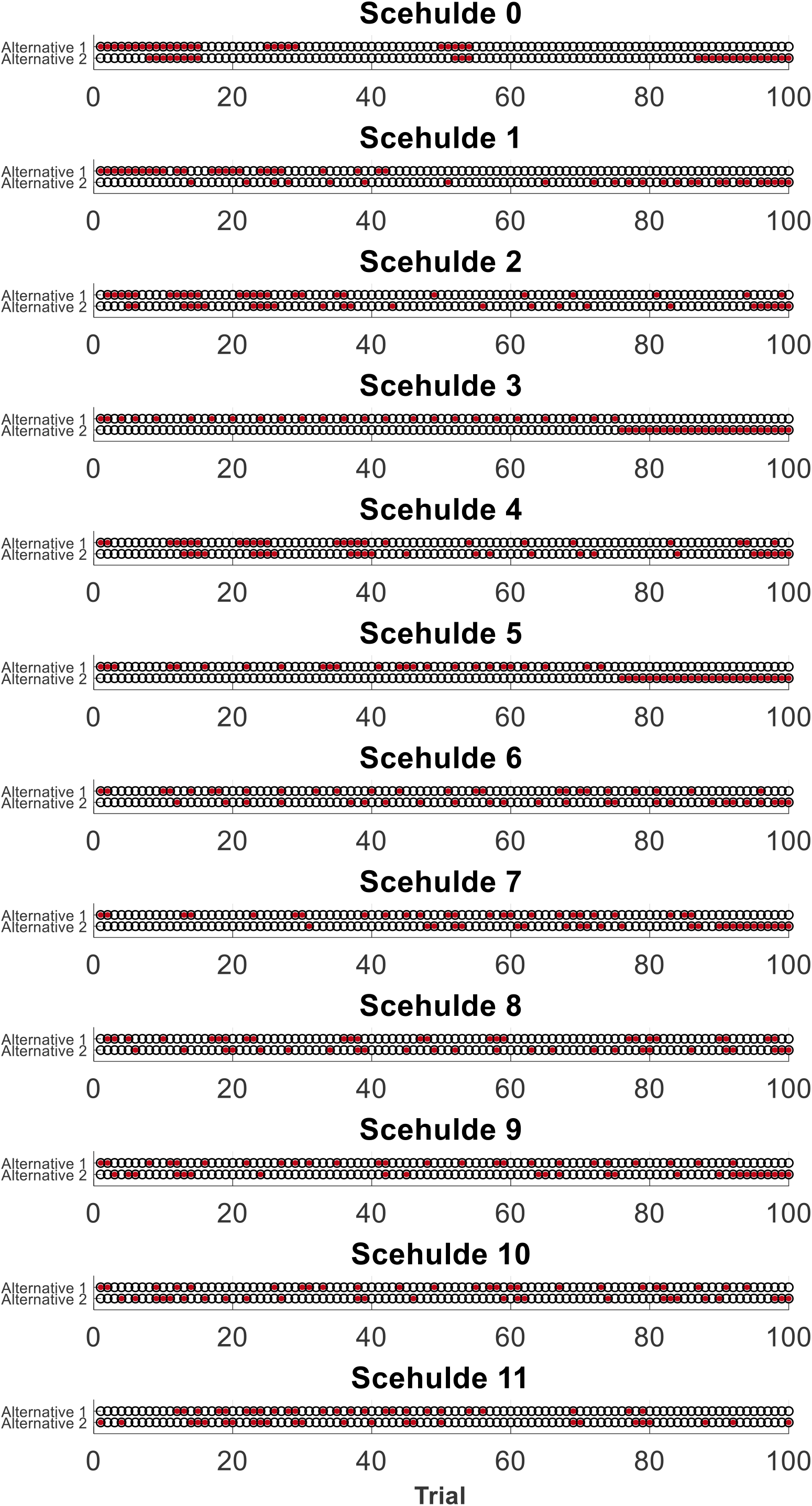
The reward schedules that were tested on human participants. Schedule 0 corresponds to the experimentally-optimized schedule. Other schedule indexes correspond to Table S1.

**Fig S3.**
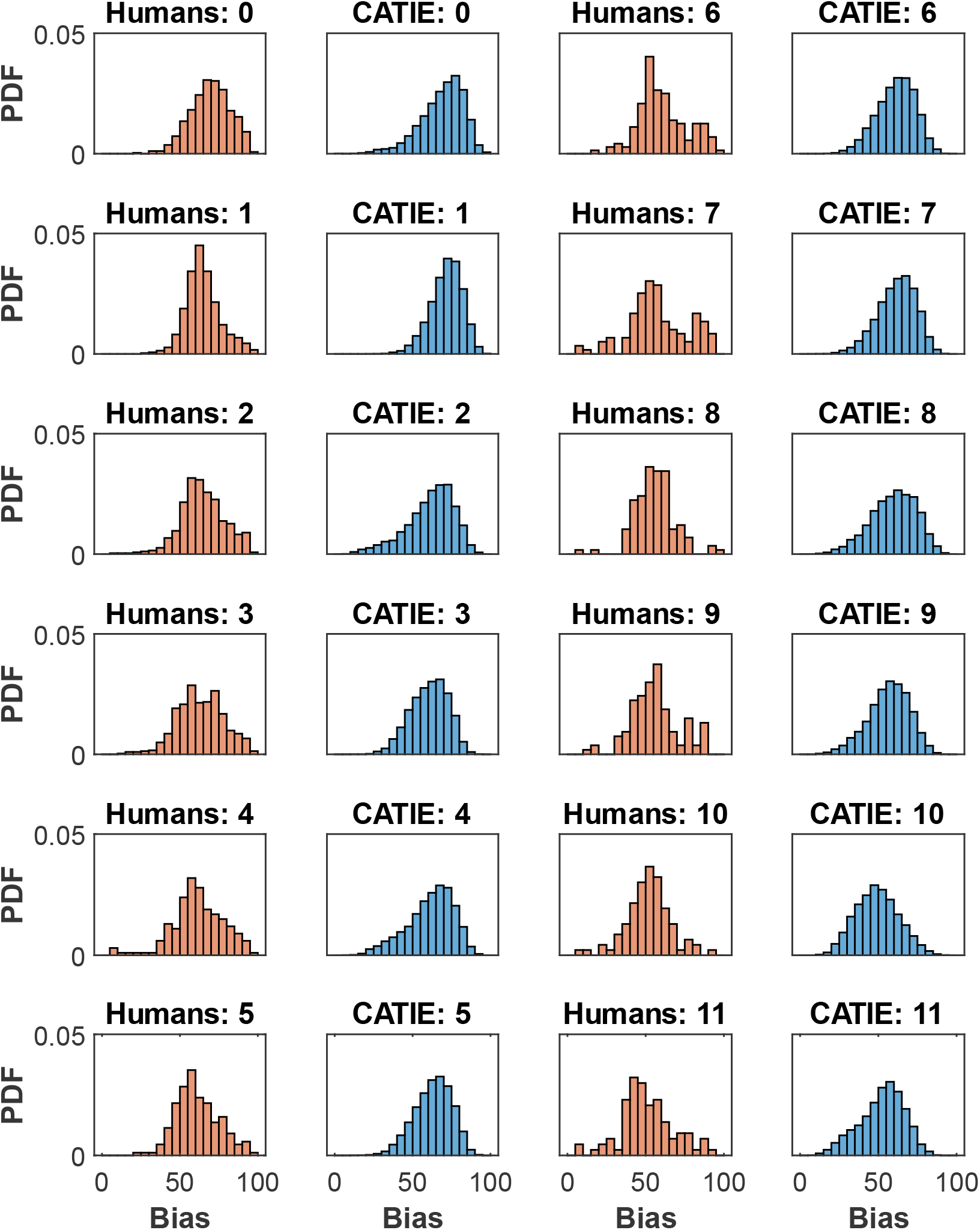
Distributions of choice biases. Red, the distributions of biases of the human subjects for each of the reward schedules in Fig. S1. Blue, the distributions of the biases of 10,000 simulations of the CATIE model for the same reward schedules.

**Fig S4.**
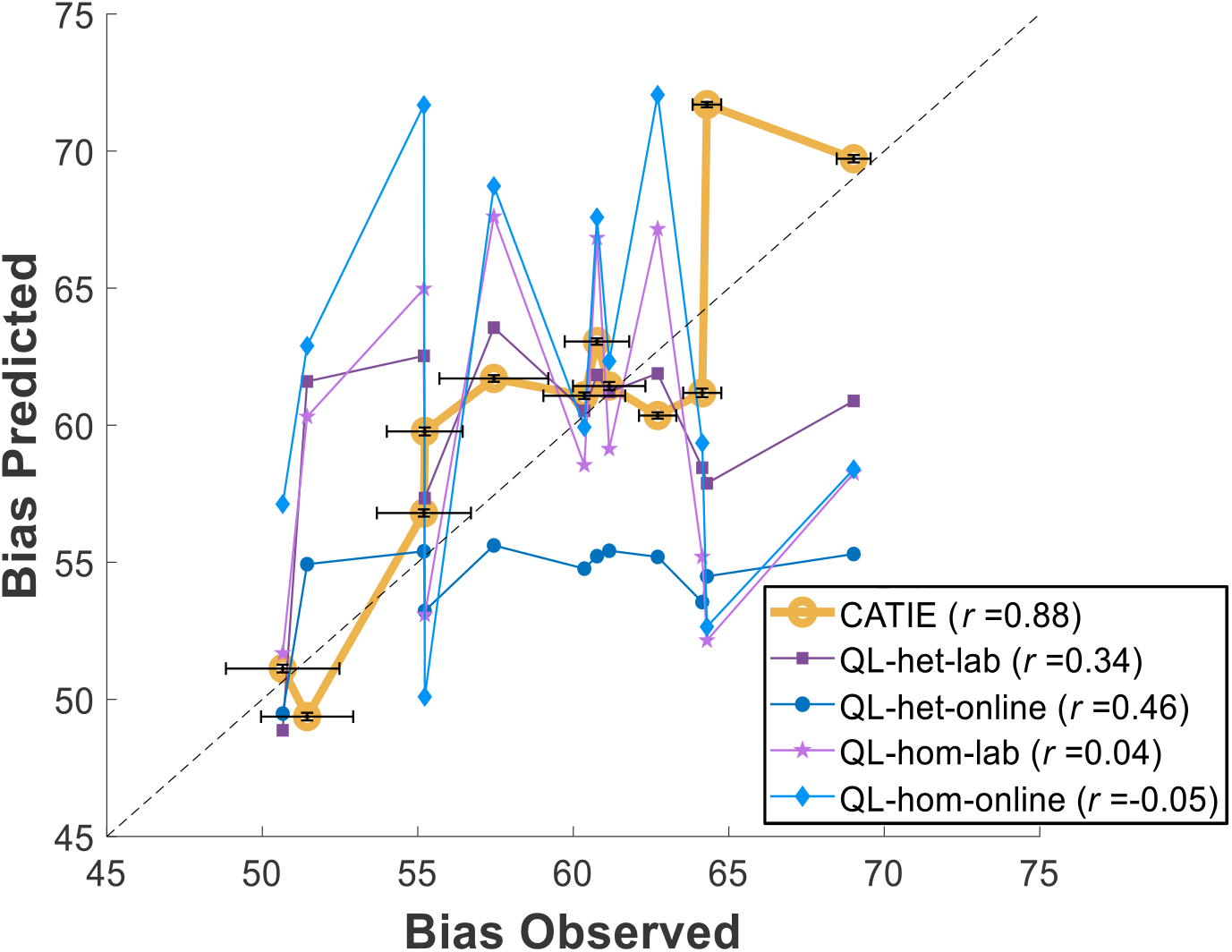
Correlation of behavioral bias with simulated bias. Both axes present the average number of choices in the target side (the “bias”) for the 11 schedules tested in the competition, with the 1 additional ideal-engineer schedule (see Results). The abscissa presents the sorted average bias as measured from behavior of the competition’s experimental participants. The ordinate presents the average bias for the same reward schedules as simulated by the 5 different quantitative models. The values of *r* in the legend depict the correlation coefficient of the predicted bias and empirical bias for each of the models. Error bars are standard error of the mean. For clarity, they are only depicted for the CATIE model. As shown by the figure, CATIE has the highest correlation with the empirical bias (*r*=0.88). The CATIE results are identical to those depicted in Fig. 3a.

**Table S2.** The dynamic track of the competition. Data were collected from 695 subjects, and they were tested on six dynamic schedules. We excluded from the analysis 19 subjects (2.7%) that chose one of the alternatives less than 5 times and are therefore likely to have ignored the reward schedule. Adding them to the analysis does not change the ranking of the schedules and their choices are posted together with those of the included participants. We did not collect any demographic information about the subjects. The number of possible dynamic schedules is far larger than that of static schedules (about 10^1500^ larger). Therefore, the ability to bias choices with a dynamic schedule is far larger than this ability with a static schedule. Indeed, the dynamic schedules were more effective than the static ones. Regrettably, only choice architects participated in the dynamic competition and therefore, we could not use it to compare choice architects to choice engineers. The winning dynamic schedule was proposed by Oren Amsalem and Haran Shani.

| Model ID | N | Bias | Model Name | Model description |
| --- | --- | --- | --- | --- |
| 1 | 125 | 69.9<br>±1.4 | Dynamic carrot and stick | The model rewards the subject for an increasing number of consecutive choices of the biased side. It is adaptive (The gap between 1 prize to another increases with time and achievements) and, avoids rewarding the unbiased side initially, in order to relate this side with failure and the other side with success. Every time the subject chooses the unbiased side, his counting is zeroed, and he needs to gain the consecutive number of choices of the biased side once again from scratch, in order to receive its reward. In order to avoid frustration and to tempt subjects to choose the biased side even in sessions where the subject did not reveal the logic of the model, after a long enough time without rewards, the biased side will be rewarded with no further conditions, in order to strengthen the association between this side and reward. We postulate when a subject will feel that he understood the pattern he will stop searching and will persist in clicking at the biased square.<br>The model calculate at each step the number of prizes that it can give, and will set the step sizes accordingly. |
| 2 | 128 | 69.4<br>±1.4 | Carrot and stick | The dynamic model contains the following rule - which will maximize the choices towards the biased side:<br>The model rewards the subject for an increasing number of consecutive choices of the biased side. It is adaptive (The gap between 1 prize to another<br>increases with time and achievements) and, avoids rewarding the unbiased side initially, in order to relate this side with failure and the other side with success. Every time the subject chooses the unbiased side, his counting is zeroed, and he needs to gain the consecutive number of choices of the biased side once again from scratch, in order to receive its reward. In order to avoid frustration and to tempt subjects to choose the biased side even in sessions where the subject did not reveal the logic of the model, after a long enough time without rewards, the biased side will be rewarded with no further conditions, in order to strengthen the association between this side and reward. We postulate when a subject will feel that he understood the pattern he will stop searching and will persist in clicking at the biased square |
| 3 | 140 | 61.9<br>±1.2 | Increasing delay contingent rewards | The model is based on the idea that agents tend to assume (explicitly or implicitly) that there is a pattern to uncover in such tasks (see Plonsky et al., 2015; Plonsky & Erev, 2017; Plonsky & Teodorescu, 2020). To get an agent who assumes a pattern exists to choose the target alternative, the pattern used here is of rewards contingent on number of consecutive choices of the target alternative. That is, to get a reward (from the target), the agent needs to choose it a number of consecutive times. In order to increase the chances that the agent captures the underlying pattern, while maintaining a reasonable delay so that rewards are not exhausted early on, this "delayed contingent reward" is provided with increasing delay times: In the first several trials, there is no delay (each choice of the target is rewarded), then reward is given only every 2nd consecutive choice of the target, then every 3rd etc. The change of delay times is contingent on the behavior of the agent (or when too many rewards were wasted): If the agent chooses in a manner that suggests she learnt the rule, the delay is increased. Moreover, allocation of rewards to the anti-target are also using the idea of pattern search. The allocation is made (starting trial 41 to make sure the agent learns early to prefer the target) every 4th trial, deterministically (based on the idea that deterministic patterns are easier to follow, see Plonsky & Teodorescu, 2020). Scheduling is thus based on the assumption that after capturing the pattern, the agent will prefer to choose 3 times the target (which is sometimes rewarded in these times) and only once the anti-target. To make sure all rewards are allocated, the length of the pattern decreases towards the end of the task. |
| 4 | 142 | 60.2<br>±1.1 | Trst | Partial reinforcement, interval prolonging with practice |
| 5 | 79 | 57.0<br>±1.3 | Pattern<br>based<br>decision | The model builds on three main observations. The first is the study of shaping (Skinner, 1953) that demonstrates the value of high initial probability of reinforcement. Second is the suggestion that learners try to detect patterns (see e.g., Plonsky & Erev, 2017). The third is the finding that surprise-triggers-change (Nevo & Erev, 2012).<br>Based on these observations our model starts with high probability of reward:<br>100% for the first biased choice, and then after each of the next 5 pairs of biased choices.<br>Next, the reward for will be given after each set of 4 Biased choices. To keep fixed pattern and reduce surprises, this schedule will be kept as long as possible. |
| 6 | 62 | 56.1<br>±1.8 | Pattern-<br>based<br>decisions 3 | The model builds on three main observations. The first is the study of shaping (Skinner, 1953) that demonstrates the value of high initial probability of reinforcement. Second is the suggestion that learners try to detect patterns (see e.g., Plonsky & Erev, 2017). The third is the finding that surprise-triggers-change (Nevo & Erev, 2012).<br>Based on these observations our model starts with high probability of reward:<br>100% for the first biased choice.<br>Next, the reward for will be given after each set of 3 Biased choices. To keep fixed pattern and reduce surprises, this schedule will be kept as long as possible. |

### Supplementary text – Sample size

To identify the most effective schedule, we utilized the method of Successive Rejects (Audibert, Bubeck, & Munos, 2010). According to this method, more subjects (more “samples”) are allocated to better-performing reward schedules. The method of Successive Rejects operates iteratively. In the first iteration, all reward schedules are allocated an equal number of subjects. Then, the worst performing schedule (the schedule associated with the minimal bias) is “rejected” in the sense that no more subjects are allocated to it. The process continues iteratively until just two schedules remain. The schedule that yields the highest bias over all tested schedules is considered the most effective one. According to this process, the number of subjects that should be used in each iteration depends on (1) the total number of schedules tested, (2) the (unknown) distribution of their “true” biases and (3) the desired confidence level for identifying the most effective one. We pre-determined the number of subjects per iteration assuming the true bias of the schedules is evenly distributed between 50% (chance level) and 70%. One minor complication when running online experiments is that not all the subjects who start the task also end up completing it. We randomly allocated subjects to schedules in advance, which resulted in some variability in the number of subjects that complete the task per schedule. To make sure that each schedule is tested on no-fewer than the number of subjects required by the Successive Reject method, based on pilot data, we pre-allocated 30% more subjects than required to each schedule.

The data presented in the paper was collected in three phases:

Phase 1: In the formal part of the competition, we used the method of Successive Rejects with a confidence level of 95%, and tested 10 of the schedules (0-6, 8-9, and 11, see below) on 941 participants. We found that the ideal engineer (schedule 0) was the most effective schedule, where schedule 1 being the second most effective schedule. We declared schedule 1 to be the winner of the competition.

Phase 2: After concluding the competition, we decided to test two additional schedules that were not included in the formal part of the competition (schedules 7 and 10). To do that, we ran a pseudo-competition over all 12 schedules using the same Successive Reject method, this time requiring confidence level of 99%. A higher confidence level entails a larger number of subjects per iteration, and a larger number of schedules entails a larger number of iterations. We repeated the process of Successive Rejects for the pseudo-competition. The initial subjects used in the earlier iterations were those collected in the formal part of the competition, and when needed we collected additional subjects. We concluded this pseudo-competition with 2,633 subjects (941 of those collected in the formal competition). The results of the pseudo-competition were similar to those of the competition: the ideal engineer (schedule 0) was the most effective schedule, followed by schedule 1.

Phase 3: By the end of phase 2, the bias induced by schedules 1-3 was rather similar. We were wondering if by increasing the number of subjects, we will be able to better distinguish between them statistically. We added subjects as to equalize to number of subjects per schedule (as before, we allocated more subjects than necessary, by a factor of 1.3, because of dropout). This increased the total number of subjects to 3,332. The ranking of the schedules was not different from that of Phase 2.

